# Effects of biological fixation methods on stimulated Raman scattering microscopy signal

**DOI:** 10.1101/2025.11.11.687741

**Authors:** Chiara Ceconello, Wuji Han, Bryce Manifold, Dario Polli, Aaron Streets

## Abstract

Stimulated Raman Scattering (SRS) microscopy enables label-free imaging of cells and tissues in their native state with chemical specificity. However, there are often experimental advantages of chemical fixation of samples prior to imaging, which can introduce perturbations that may alter the native state of the samples, and possibly impact the SRS signal. In this study, we systematically characterize the effects of common fixatives (Paraformaldehyde, Formalin, Glutaraldehyde, Methanol, Ethanol) on the preservation of cellular integrity and molecular composition in a Hela cell model as observed by SRS microscopy. We demonstrate how the different fixatives can influence lipid and protein content, and overall cell morphology, with significant implications for the accuracy of quantitative SRS microscopy. Our findings indicate that Paraformaldehyde (PFA) imposes minimal disruption to cellular and molecular states compared to the other fixatives tested, and suggest Glutaraldehyde (GA) as a suitable alternative for SRS imaging. This study provides insights for the choice of the optimal sample preparation, enabling more reliable SRS-based studies for the evaluation of cellular processes and disease mechanisms where fixation is used.

## Introduction

Stimulated Raman Scattering (SRS) is a powerful microscopy technique for label-free investigation of cells and tissues by exciting distinct molecular vibrations to provide unique endogenous signatures for chemical profiling^1,2^. Offering high-speed quantitative imaging, with high spatial resolution in three dimensions, SRS has demonstrated great potential in metabolic studies and tissue diagnosis^3,4^ without the need for chemical labeling. This allows unperturbed characterization of biological samples, which can be critical to accurately understand biological function in situ. However, while chemical labeling is not required, many SRS studies are performed on fixed samples, calling for preparation protocols to preserve sample characteristics as close to the native state as possible. Chemical fixation, developed in the late nineteenth century for preserving tissues for microscopic examination^5,6^, involves adding solvents or cross-linking molecules to samples to “freeze” intracellular molecular content, preventing post-mortem decay and providing additional mechanical strength and protection against bacteria^5,7^. As fixation alters molecular structures, a precise, sensitive, and comprehensive characterization of biomolecular changes is critical for accurate interpretation of SRS images when using fixatives^8^. Moreover, in experiments requiring the integration of diverse analytical techniques for phenotypic characterization (e.g. SRS microscopy, mass spectrometry, or transcriptomics), understanding compatible fixation methods and their specific effects in cellular chemistry and morphology, ensures clear and accurate results. Vibrational spectroscopy itself, being inherently label-free and chemically specific, is an excellent method for analyzing the effects of fixation on cells^9–13^. Spontaneous Raman spectroscopy has been used to investigate the effects of sample fixation across various bacteria strains^14^, eukaryotic tissues^14,15^, and different human cell lines^13,16–18^. These fixation-induced changes are characterized spectrally through analysis of key peaks in the fingerprint region of the vibrational spectrum.While spontaneous Raman spectroscopy provides valuable molecular information, its low signal levels make it impractical for fast, high-resolution imaging of cells and tissues. Coherent Raman, particularly in the CH stretching region, overcomes this limitation by enabling rapid, label-free chemical imaging with high spatial resolution. Techniques like CARS and SRS have proven effective in visualizing cellular content and processes, successfully detecting intracellular lipids, proteins, nucleic acids, and water with high speed and sensitivity^3,19–23^. Despite the increasing adoption of coherent Raman imaging in biological research^24–26^, a systematic characterization of how fixation affects these measurements is still lacking. In this work, we address this gap by exploring the effects of common fixatives on SRS microscopy images of biological samples, providing a more comprehensive characterization of fixation-induced changes through high-resolution chemical and morphological features.

Here, we explore the effects of common fixatives on SRS microscopy images of biological samples (Fig. 1). We offer insights into both the morphological and chemical effects of these fixatives on HeLa cells, a well-characterized, widely-used, and reliable cell line model that serves as an excellent platform for exploring fixation effects. We extend the analysis of cell fixation via a broad range of protocols explored in the CH region through SRS microspectroscopy, providing additional quantitative morphological insights. We select five commonly employed fixatives, chosen for their widespread adoption in microscopy and their popularity in adjacent fields (e.g. DNA, RNA sequencing) to evaluate against live samples. This study offers a broader characterization of fixation-induced changes in SRS microscopy, aiming to inform the design of experiments that integrate label-free chemical imaging with orthogonal approaches.

**Figure 1:**
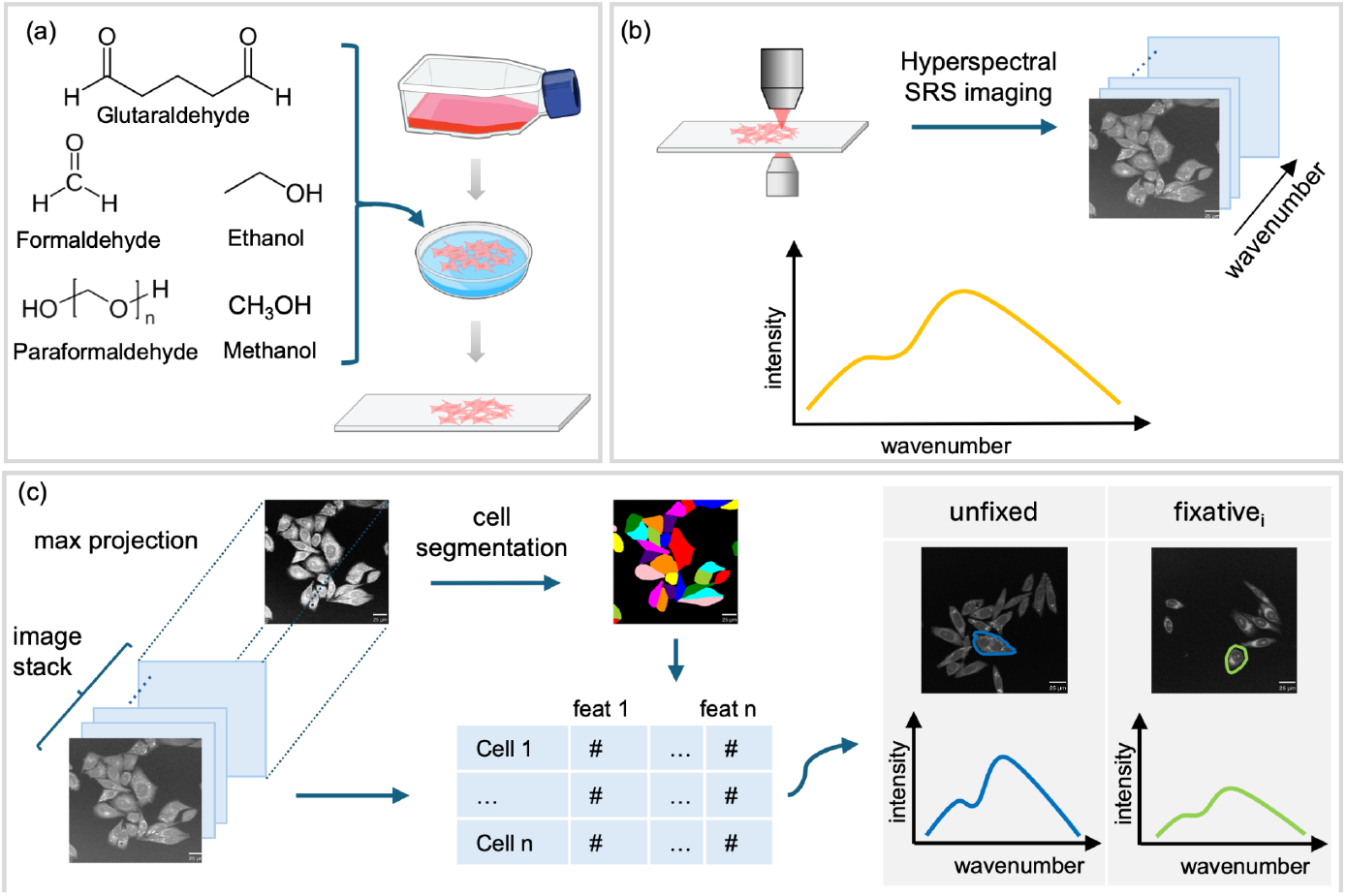
Schematic representation of the fixation SRS experiment. (a) Cell culture, fixation and seeding on microscope slides. (b) Stimulated Raman Scattering data hypercube acquisition in the CH spectral region. (c) Image post processing workflow: each hyperstack is pre-processed and segmented via cellpose^48^ to extract single-cell metrics and build a comprehensive Raman-shift specific data frame. Then fixatives are compared to the live counterpart both spatially and spectrally.

We aim to provide supporting evidence that PFA (or other cross-linkers, e.g. NBF) is indeed the least disruptive and recommended choice for coherent Raman microscopy-based assays. We also evaluate the effects of glutaraldehyde and propose it as a suitable alternative for SRS imaging, owing to a potentially non-disruptive signal enhancement^27^. Finally, we quantify the effects of lipid loss and cell shrinkage resulting from alcohol-based fixation.

Commonly used fixatives include aldehydes (formaldehyde and glutaraldehyde), alcohols (methanol, ethanol), ketones (acetone), and acids (tannic acid, picric acid, etc.). When used individually, these fixatives often induce undesirable chemical alterations in biological samples, and it is common to use them in combination to balance out unwanted effects^17^. The predominant cell fixative employed in biology is formaldehyde, typically used in a polymer form such as Paraformaldehyde (PFA) at 4% dilution, in an aqueous solution of formaldehyde gas (37-40% by weight) with a small amount of methanol added to prevent polymerization, or as neutral buffered formalin (NBF) at 10% dilution. It is highly used in SRS microscopy^24,28–30^, as formaldehyde-based fixatives are thought to preserve major biochemical components of the cell in a form that is closest to that in the live cell^17^. During fixation, the methylene glycol component of formaldehyde forms cross-linkages between the NH_2_ terminals of protein amino acid residues and the NH group of a tertiary amide^11^. Fixation in formalin thus adjusts the vibrational freedom of membrane lipids, possibly through a reaction of methylene glycol in aqueous formalin with unsaturated hydrocarbon chains^9^, causing lipid leaching^31^. Formaldehyde fixation also disrupts hydrogen bonds within large intracellular molecules^7^ and denatures intracellular proteins by forming cross-links among amine residues ^7,9,32^, leading to the loss of quaternary, tertiary, and possibly secondary structures of proteins^33^. Formaldehyde fixation has also recently been shown to change intracellular hydrogen bonding spectra in SRS microscopy, suggesting a significant change in free intracellular water^34^. Protein cross-linking may cause overall sample shrinkage^7^ and formation of nanoscale structures caused by aggregation of membrane proteins both in formaldehyde and glutaraldehyde fixation^35^. Moreover, prolonged exposure to formaldehyde reduces antigen binding specificity in immunohistochemical staining^33^, decreases DNA solubility, and slowly hydrolyzes the DNA and RNA backbones^7,36–38^ effectively reducing nucleolar RNA concentration^8^. Other previous work has shown formalin fixation unstacks DNA bases in adenine-thymine-rich regions, forms hydroxyl-methyl groups with nucleic acid bases, and hydrolyzes N-glycosidic bonds, resulting in free purine and pyrimidine groups^7^. Due to the loss of nuclear integrity^13^, alcohols are generally preferred to formalin for their better preservation of antigens, despite causing overall cell shrinkage^39,40^. Based on this, we expect formalin-fixed cells to appear smaller when imaging with SRS, due to overall shrinkage from intracellular water loss and to show a potential reduction both in the CH_2_ and CH_3_ peak due to lipid leaching and disruptions to protein folding and structure.

Glutaraldehyde, typically used in 1.25-2.5% dilutions, is an alternative to formaldehyde-based fixation. It forms covalent cross-links between amine residues in proteins^14,17^. Compared to formaldehyde, glutaraldehyde penetrates cells and tissues more slowly but exhibits stronger cross-linking due to its two aldehyde groups^31^. This makes it particularly suitable for electron microscopy, where stronger cross-linkers are preferred^41,42^. Additionally, like formalin-based fixatives, glutaraldehyde preserves cytoplasmic lipid content^9^ and has been observed via Raman spectroscopy to increase CH_2_ bending and CH_2_/CH_3_ twisting relative to live cases^27^. Due to its stronger cross-linking causing higher structural rigidity, we expect effects from glutaraldehyde to be less pronounced than formaldehyde in size variation. However, a more pronounced change in CH_2_ and CH_3_ signals could be observed potentially reflecting closer-to-live levels or higher intensity due to significant structural and chemical rearrangements.

Finally, alcohols such as methanol and ethanol are thought to better preserve nucleic acids but have strong denaturing effects on proteins^31^. They may solubilize membrane lipids^43^, causing lipid leaching^44,45^ and overall tissue shrinkage^8^, which is why they are often combined with tissue-swelling fixatives. Despite causing protein rearrangement favoring β-sheet over α-helical conformations ^46^, the cell shrinkage effect increases concentrations of nucleolar proteins and RNA, making alcohol fixation preferred for RNA based assays such as transcriptomics^8^. In single-cell transcriptomics, methanol-fixed samples maintain a cellular composition similar to fresh samples, providing good cell quality with minimal expression biases^47^. The precipitation of other macromolecules (i.e., aggregation out of solution) occurs over several minutes^35^, with denatured proteins partially refolding upon rehydration in Phosphate-buffered Saline (PBS)^46^. We expect SRS images of alcohol-fixed cells to show little to no lipid signal, significant shrinkage due to dehydration, and an overall reduction in protein signal, with higher protein concentration localized in the nuclear region.

## Methods

### HeLa Sample Preparation

HeLa cells (UC Berkeley Tissue Culture Facility) were cultivated and maintained in Dulbecco’s Modified Eagle’s Medium (DMEM) (Corning) supplemented with 10% Fetal Bovine Serum (FBS) and 1% PenStrep (Penicillin, Streptomycin, L-glutamine) at 37°C and 5% CO_2_. In preparation for imaging, cells were plated at 20-30% confluence on 8.8cm^2^ petri dishes (Nunclon Delta Surface, Thermo Scientific), each prepared with a ∼0.15mm-thick micro cover glass (VWR), and allowed 24 hours to adhere.

At ∼50% confluence, cells were washed three times with 1X PBS (Corning). For live imaging, the cells plated on a micro cover glass were covered with a microscope slide (VWR Vistavision, glass 1-mm thick) to create a sandwich-like configuration with 20 μL of 1X PBS to prevent drying. Multiple fields of view were acquired within 30-40 minutes of removal from media to avoid live cell alteration due to drying.

### Fixation protocols and experiment design

For cell fixation (Fig. 1(a)), five mixtures were prepared as:

1. A 4% Paraformaldehyde (PFA) solution, pH balanced in PBS (Pre-mixed, Invitrogen Thermo Fisher), stored at 4°C;
2. 10% Neutral Buffered Formalin (NBF) in PBS (Pre-mixed, G-Biosciences), stored at 4°C;
3. A 1.5% Glutaraldehyde (GA) (25% initial aqueous solution, Thermo Scientific) solution, pH balanced in PBS, stored at 4°C;
4. A 10% Methanol (MetOH) (99.8%, Thermo Scientific) dilution in molecular-grade water, stored at -20°C;
5. A 70% Ethanol (EtOH) (200 proof, molecular-grade, Fisher BioReagents) dilution in molecular-grade water, stored at -20°C;

Raman spectra of the fixation mixtures in the above percentages are provided in the supplementary material (Fig S1).

Similar to the live cell protocol, 3x washes were performed with PBS. 2 mL of mixture were added to the petri dish and allowed to fix for 15 minutes at room temperature for cross-linking fixation (PFA, NBF, GA) or 10 minutes at -20°C for alcohols (MetOH, EtOH). In all protocols, mixtures were then aspirated and followed by 3x washes with PBS. Fixed cells on the micro cover glass were then sandwiched with a microscope slide with 20 μL of PBS to prevent drying, and imaged.

For each live/fixed condition, we prepared two sample replicates, and multiple fields of view were acquired to minimize sample-dependent effects. This experimental workflow was replicated on different days to control for sample-dependent and day-dependent variations.

### Stimulated Raman scattering microscope

A dual-output Insight DS+ laser operating at 80-MHz repetition rate, emits 100-200f pulses at the output: a 1040nm Stokes beam and tunable pump set at 799nm, targeting the CH spectral region, covering the 2820-3100 cm^−1^ range through spectral focusing SRS^49,50^. Stimulated Raman loss (SRL), is measured modulating the Stokes beam through an EOM (Thorlabs) at 10.28 MHz. The Stokes power is controlled via a motorized half-wave plate-polarizing beam splitter pair. Both beams are chirped with three H-ZF52 glass rods (Union Optic), reaching ∼2 ps duration. A motorized linear delay stage (Newport FCL200) and broadband retro reflective mirror (Newport) pair, adjusts pump-Stokes beam overlap to achieve spectral focusing in the CH region. A dichroic mirror recombines pump and Stokes beam, then coupled with a fully automated commercial inverted microscope (Olympus IX83). Both excitation beams are focused on the sample, mounted on a motorized XY stage (HLD117 Prior), with a water immersion 60x, NA=1.2 objective (W/IR UplansApo, FN26.5, Olympus) and collection is via an oil-immersion condenser with NA=1.4 (D-cuo achr-apl Nikon - CSC1003, Thorlabs). The SRL signal is retrieved demodulating the pump beam with a Lock-in amplifier (HF2LI Zurich Instruments) after being filtered (FESH1000 Thorlabs) and collected with a photodiode (Hamamatsu S3590-08). Prior to imaging, the system is calibrated each day using 20-50 μL of 25% DMSO sandwiched between a microscope slide and coverslip, and a single-point spectrum is acquired to match the delay line position with known spectral peaks. For cell imaging, powers are set to 20/25 mW (Pump/Stokes). For each measurement, we acquired hyperspectral images at 16 spectral points, every 12.62 cm^−1^ corresponding to Raman shifts ranging from 2850 to 2920 cm^−1^, targeting both lipid and protein peaks (Fig. 1(b)). Each acquired field of view (FoV) is 512 by 512 pixels, covering 212 x 212 μm^2^. To ensure consistency across different imaging days, all measurements were corrected to the highest lock-in gain. Prior to analysis, the raw image hyperstacks underwent field correction by normalization with a 25% DMSO calibration solvent image. To be specific, in each experiment, a Gaussian filter with a sigma value of 50 (FWHM = 117.74 pixels to match the DMSO image) was applied on the average of three DMSO solvent images, and divided by its maximum value. This broadband filter was used to reduce acquisition noise due to the inability to take paired dark pixels at each position, and as a consequence the field correction was relative instead of absolute. The pixel-wise inverse of the reference solvent image was used as a normalizer on cell sample images in the same experiment. This procedure enhances signal uniformity, particularly at the edges of images, which are typically less intense due to aberrations introduced by point-scanning (FV12-SU high sensitivity galvo scanning unit, part of Olympus FV1200 microscope system) through the high-NA objective across the field of view.

### Single Cell Segmentation and Statistics

To isolate single cell spectra from the images, we used the pre-trained `cyto3` model from Cellpose^48^ to perform cell segmentation. A maximum projection of the hyperspectral image stack was first performed to create a 2D image for Cellpose. Then, for each image, a U-Net is used to generate horizontal and vertical gradients, which forms a vector field. By grouping together pixels that converge to the same center, individual cell masks could be drawn.

After creating the mask image from the output, individual cell labels are used to iterate through summary statistic calculations. For example, the sum of intensities adds up all the pixel values from the same cell, and the *k*-th *q*-quantile calculation outputs the intensity value based on the cumulative distribution of the intensities.

For each field of view, a table with cell labels as rows and statistical type as columns can be generated. To merge multiple fields of views and experiments, each image gets a unique label based on the concatenation of the experiment label, field of view label, and cell label.

### Spectral Similarity Examination via Cross Correlation

Given that spectra across conditions are near-identical, an unbiased way to compare spectral shape is preferred. This is achieved by calculating correlations between normalized spectra. The bootstrapping test for each condition pair is designed as follows.

First, 200 cells are sampled from either condition. The spectrum for each cell based on average intensities at different Raman shifts were extracted. These spectra were then normalized by dividing its maximum value along the spectral axis (Raman shift), in order to average them into one single spectrum.

A pair of average spectra were used for each condition pair, to calculate Pearson product-moment correlation coefficient, i.e.

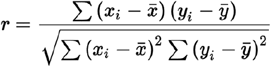

The sampling is then repeated for a total of 100 times. This would yield the mean and standard deviation of *r*.

Since *r* is very close to 1, the log scale representation was used for visualization, where *s*=-log(1-*r*) is used to indicate how close *r* is from 1. Thus, the closer *r* is to 1, the higher the similarity (*s*) between spectra.

### Spectral deconvolution through N-FINDR

We isolated component spectra contributions through unmixing via the N-FINDR algorithm^26,51^. The dimensionality of the hyperspectral image is reduced by finding *n* pure spectral components that best represents the data. To achieve that, the N-FINDR algorithm optimizes the volume of a s-dimensional simplex in a (s+1)-dimensional space, here s denotes the spectral dimension. The vertices, i.e. endmembers of the simplex become the “pure spectra” after the optimization, and their relative abundance in each pixel can be reconstructed throughout the whole image, utilizing methods such as non-negative least squares fitting.

## Results

### Single-cell analysis reveals fixation-dependent variation in Raman spectra

We used SRS microscopy to image the fixation conditions described in Table 1. For each fixative, we acquired a set of fields of view from replicates across multiple days. We performed a max projection across the wavenumber axis to provide an estimate of overall signal intensity and used these images to visualize the impact of fixation on cells (Fig. 2 (a-f)). With respect to unfixed cells, we observed higher overall SRS signal for glutaraldehyde fixed cells, some signal reduction in paraformaldehyde and formalin fixed cells, and a more prominent signal change for alcohols.

**Table 1:** Summary of Cell Counts and Replicates for Hyperspectral SRS Imaging Across Fixation Conditions.

| Fixative | Replicates | Number of Cells |
| --- | --- | --- |
| 1.5% glutaraldehyde | 6 | 327 |
| 10% formalin | 6 | 348 |
| 4% PFA | 6 | 308 |
| 70% ethanol | 6 | 331 |
| 90% methanol | 6 | 407 |
| unfixed | 10 | 604 |

**Figure 2:**
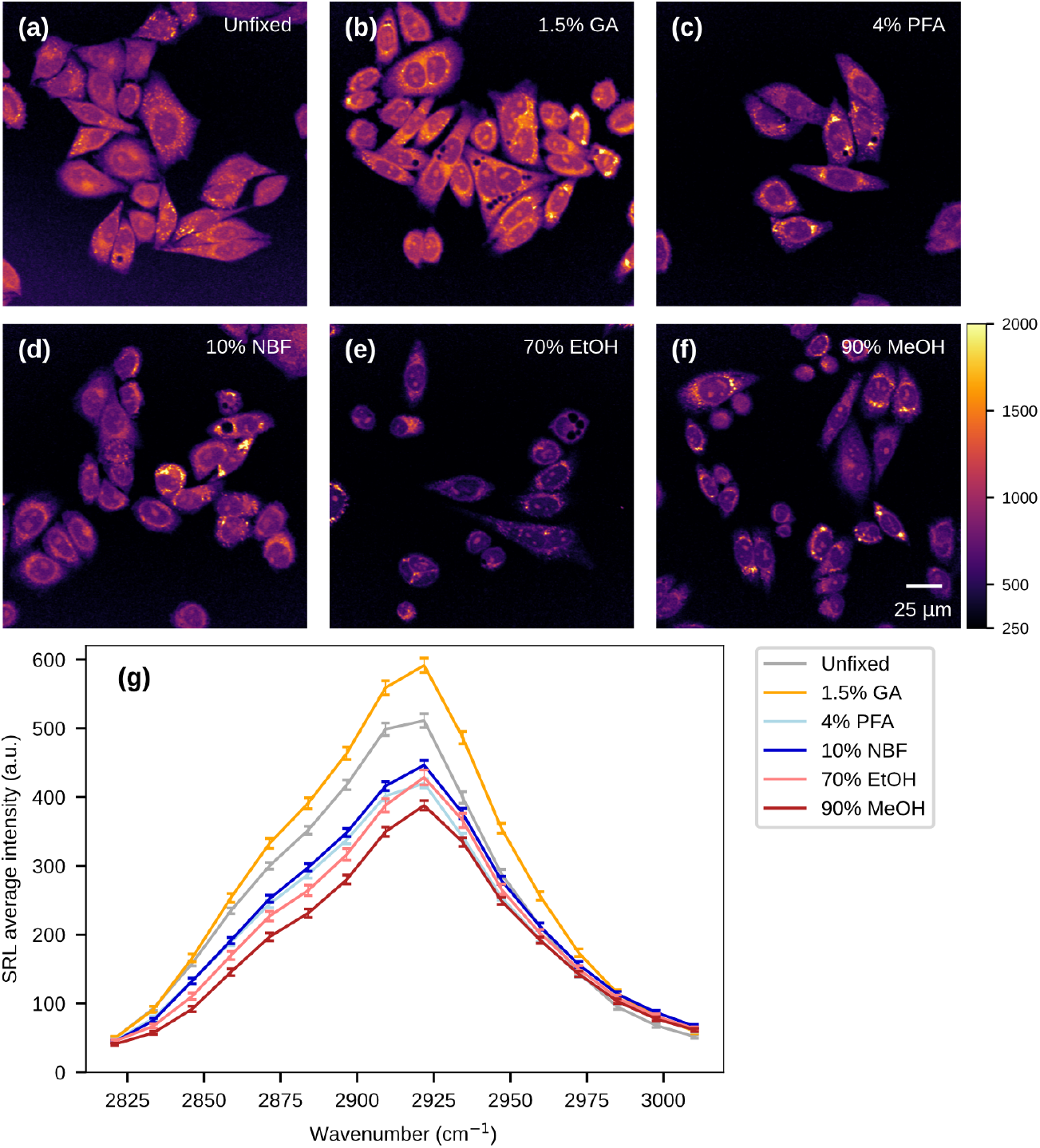
SRS hyperspectral images of fixed and live cells reveal morphological and spectral differences across conditions. (a-f) Max projection across Raman signal of six SRS images, one per condition, respectively: unfixed, glutaraldehyde, paraformaldehyde, formalin, methanol, ethanol. With respect to live, glutaraldehyde shows higher overall SRS signal, paraformaldehyde and formalin a small decrease, while alcohols show the highest signal variation. The behaviour is confirmed in (g): average intensity of single cell spectra across all samples grouped by fixation condition. Vertical error bars represent 95% confidence intervals of the mean estimate.

We then explored SRSspectra of single cells across all of the conditions in Table 1. To extract single-cell SRS spectra from the hyper-spectral image stacks, we used Cellpose^{Updating}^, a pre-trained model for segmentation, to create individual cell masks for single-cell metric extraction. Individual cell masks were segmented with the maximum projection image (See methods). Each individual cell’s mask was then applied on the hyperspectral stack, creating summary statistics at every Raman shift for each cell. The summary statistics include sum of SRS intensity, area, and intensities at different pixel quantiles. We constructed a comprehensive data frame of 2325 cells (Table 1) across all experimental conditions (live, fixed), and imaging days, and extracted Raman shift-specific average signal intensity (cumulative cell intensity divided by its size) and cell size across all samples and conditions (Fig. 1(c)).

These single-cell average spectra are presented in Figure 2(g) organized by fixation condition. To minimize potential biases induced from cell-size differences, single-cell cumulative intensities are divided by their respective area. In glutaraldehyde fixed cell spectra, we observed an overall signal increase with respect to unfixed (16% increase at the 2920cm^-1^ band, see supplementary table S2), consistent with previous reports of Raman signature changes^27,52^, and our observation on max projection images that appear consistently brighter (Fig. 2(a)). Paraformaldehyde and formalin cause a signal reduction, of respectively 18% and 16% decrease at the 2920cm^-1^ band, in agreement with previous works^8,13,17^. Alcohol fixation results appeared to be most disruptive, with signal reductions up to 16% (ethanol) and 24% (methanol) at the 2920cm^-1^ band, coupled with visible cell shrinkage (Fig. 2(e,f)) and internal rearrangement, as reported in the literature^8^.

We calculated the cross-correlation between average spectra from different fixation conditions to further explore how fixatives perturb the chemical composition of the samples. Pearson’s product–moment correlation coefficient r was computed point-by-point across the spectra, and transformed as a measure of Similarity S=−log(1−r) to enhance the visibility of small deviations at high similarity values (See methods). This analysis distinguishes between uniform signal reduction and changes in specific spectral features. As shown in Fig. 3(a) (top row) all fixatives altered the single-cell CH spectral signature to some extent, with paraformaldehyde and glutaraldehyde showing the highest similarity values to unfixed cells, i.e. 5.3 and 5.2 respectively. While formalin showed a smaller signal reduction than paraformaldehyde in terms of intensity, it exhibited greater spectral distortion (S=4.5). Dissimilarity increases with alcohol fixation, with ethanol showing an S value of 3.8 and methanol 3.3, reflecting greater alteration of the spectral profile.

**Fig. 3:**
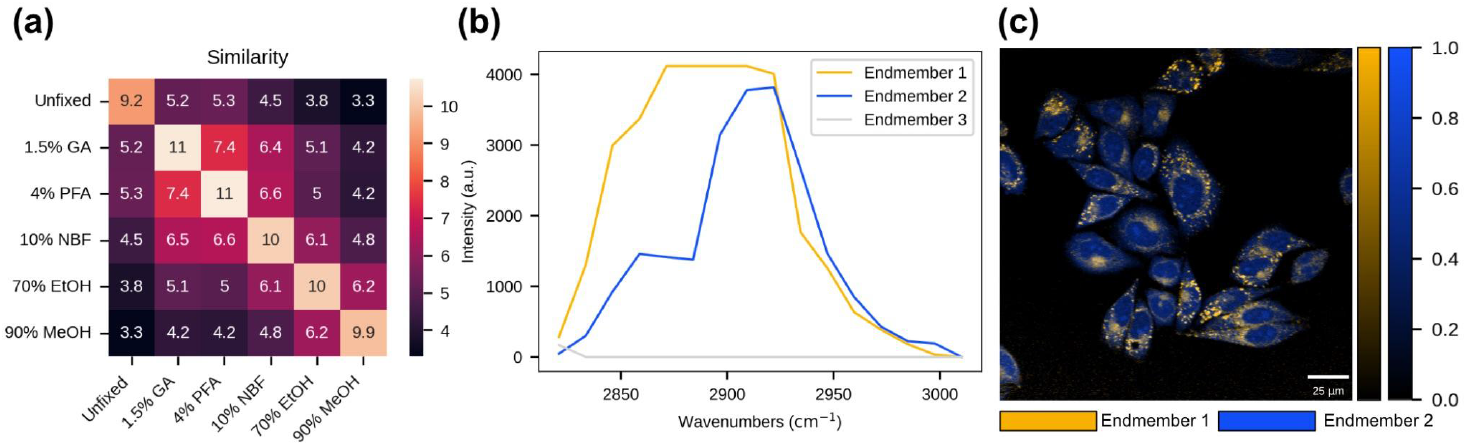
Spectral analysis on hyperspectral SRS images of cells. (a) Spectral similarity assessment by cross-correlation of spectrum pairs. Spectra from individual cells were extracted based on their average intensities at multiple wavenumbers. These spectra are then normalized by dividing their maximum spectral value. Spectra from individual cells were extracted from different fixative conditions. These spectra were then resampled per condition to perform a bootstrapping Pearson product-moment correlation coefficient calculation, between condition pairs. The similarities are calculated with -log(1-r), where r is the correlation coefficient. Heatmap shows the negative log scale, higher value indicates more spectrally similar condition pairs. (b) Endmember extraction using N-FINDR. 32 images of cells are pooled together, and applied to an n=3 N-FINDR feature extraction. A mask of cell v. no-cell area was used to eliminate background interference with the algorithm. 3 endmember spectra are found, with the last being noise. (c) Two-color reconstruction of the two significant endmembers with non-negative least squares fitting. Each endmember image has a dynamic range between 0 and 1. Scales were applied to increase visual contrast.

These observations support PFA fixation as a reliable model for live-cell representation, preserving signal intensity (with minor reductions) and spectral integrity, and identify glutaraldehyde as a strong candidate for SRS imaging in the CH region, combining signal enhancement with minimal distortion of spectral shape.

### Two-channel imaging identifies fixative-specific alteration of lipid and protein content

To better understand chemical differences induced by fixation, we isolated component spectra contributions through unmixing via the N-FINDR algorithm^26,51^. The number of endmembers, i.e. the chemical species, is selected empirically based on the number of non-arbitrary components found. Here, based on main cell constituents (protein and lipid), three components were used. We implemented this step in a reduced pixel subset from unfixed cells only, in order not to include any fixation artifacts in the spectral features. The extracted components mapped to expected biomolecular species in the CH region: lipids (Fig. 3b, endmember 1) and proteins (Fig. 3b, endmember 2). The last endmember identifies the off-cell background pixels whose pixel values fluctuate from noise (See zoom-in in supplementary Fig. S4). The extracted endmembers (Fig. 3(b)), are consistent with with Raman spectra of protein and lipid previously reported in the literature^25,53,54^. Their chemical significance was validated through two color abundance maps using non-negative least-square fitting (Fig. 3(c)). Here, the two endmembers effectively represent protein and lipid content in cells in a two-color recolored image. This full spectral decomposition justifies a lower-dimensional representation of our hyperspectral data by focusing on protein and lipid effects, choosing the spectral bands at 2850 and 2920 cm^-1^ as the contributions with the least amount of cross-talk (See supplementary Fig. S5). These bands offer high contrast for robust discrimination between lipid- and protein-rich regions and are well-established in the literature as markers for lipid^1,3,23,55^ (CH_2_ symmetric stretch) and protein^23,28^ (CH_3_ symmetric stretch) content.

Two-color composite images are generated from the hyperstacks by isolating protein and lipid signals from the CH_2_ and CH_3_ channels, respectively. The CH_2_ signal (2850 cm^−1^) corresponds mainly to methylene groups found abundantly in lipids, while the CH_3_ signal (2930 cm^−1^) indicates methyl groups, primarily associated with proteins but also present in lipids. To achieve clear separation, the pure protein signal from the CH_3_ image is extracted by subtracting a rescaled CH_2_ image, ensuring both images share the same intensity range. Six fixation conditions are visualized on a consistent color scale, to highlight the linear dependence between signal intensity and component concentration of SRS (Fig. 4 (a-f)).

**Fig 4.**
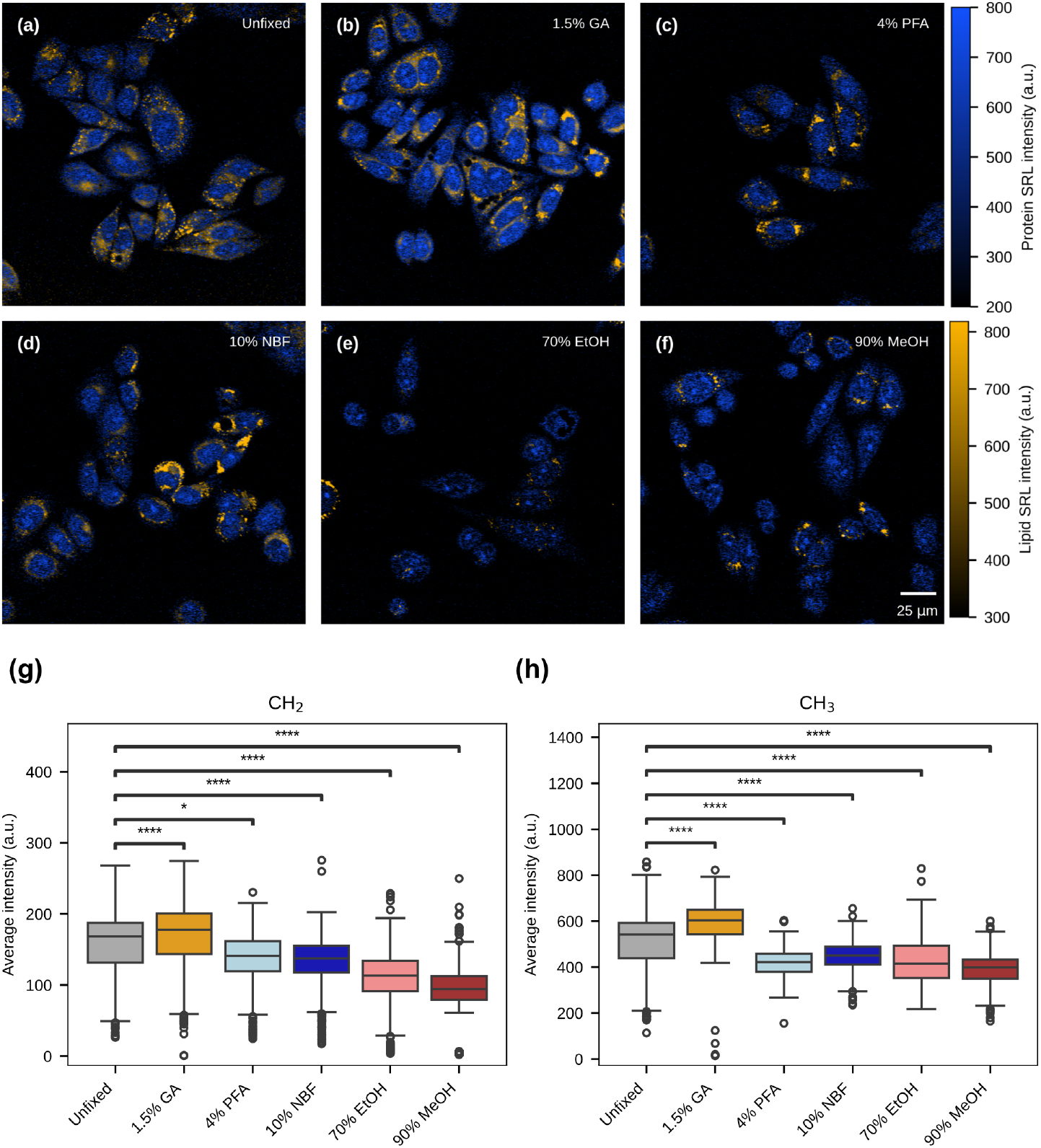
Two-color SRS images reveal fixation induced protein and lipid concentration changes on HeLa cells. (**a** to **f**) Representative two-color composite images of the lipid (2850 cm-1) and protein (2920cm-1) channels in six different sample preparation conditions, respectively: live (unfixed), glutaraldehyde, paraformaldehyde, neutral-buffered formalin, ethanol and methanol. (g,h) boxplots of protein and lipid distributions of all the cells in a specific sample preparation category. Each replicate experiment involved n>376 cells. *, **, ***, and **** indicate p < 0.05, 0.01, 0.001, and 0.0001 (Welch’s t-test), respectively. Horizontal bars of the boxes represent maximum, 0.75, 0.5, 0.25, minimum, respectively. Outliers (i.e. circles) are determined by whether the point deviates 1.5 times the interquartile range (0.75 quartile - 0.25 quartile), from either quartile.

### Cross-linking fixatives preserve protein concentration closest to live cells

Live cells (Fig. 4(a)) show bright and uniform protein signal (blue) throughout the cell. Glutaraldehyde-fixed cells, similarly show intense and uniform protein signal (blue), with signal distribution on average more intense than the live counterpart. Both PFA and NBF, with slightly fainter blue colors, show a small decrease in average protein signal and a less uniform protein arrangement, with significant structural reorganization characterized by the appearance of different bright internal nuclear structures (See Fig. S7 (c,d)). Alcohol fixation (Fig. 4(e-f)) yields the most variation compared to live cells, with an overall reduction in protein signal and a more localized signal intensity in the nuclear area where fibrillar aggregates are clearly visible (See Fig. S7 (e,f)).

### Alcohol-based fixatives cause consistent lipid loss

Live cells show bright and small lipid aggregations (gold) distributed mainly at the edges of the cell. Glutaraldehyde-fixed cells show intense lipid signal (gold), with signal distribution on average more intense than live case. Lipids in PFA-fixed cells show less significant change with respect to unfixed. Making it the most desirable method at preserving lipid content. Here, the significance values are assessed by paired Welch’s t-tests between unfixed cells and each fixative condition, on the SRL average intensity. Whilst other fixatives show a drastic difference with p-value less than 0.0001, PFA has a much higher p-value greater than 0.01. NBF, with fainter lipid signal, exhibits a higher level of washout possibly due to its methanol content (up to 2%). Alcohol fixatives most likely cause lipid solubilization and washing away^44,45^, with ethanol exhibiting some lipid loss and aggregations into larger structures with respect to unfixed. In the case of Methanol fixation, lipids are not visible and almost completely washed out.

### Glutaraldehyde fixation exhibits minimal impact on cell size

The average unfixed cell size, measured by area, is 488μm^2^ (Fig. 5), which agrees with the average cross section of HeLa cells grown in non-human serum (370±100μm^2^)^56^. Glutaraldehyde fixation has almost no impact on cell size (3.7% decrease, see table S6) (p-value<0.05). Both PFA and NBF exhibit slight size increase and swelling, possibly due to rehydration in PBS, with PFA showing the least perturbation (6.9% increase, see table S6) (p-value<0.01). In alcohol fixation, considerable shrinking can be quantified in both conditions. To support that the observed size variations across fixatives are sample independent, we evaluated size variation across different sample replicates and imaging days for unfixed cells only (Fig. S7).

**Fig. 5:**
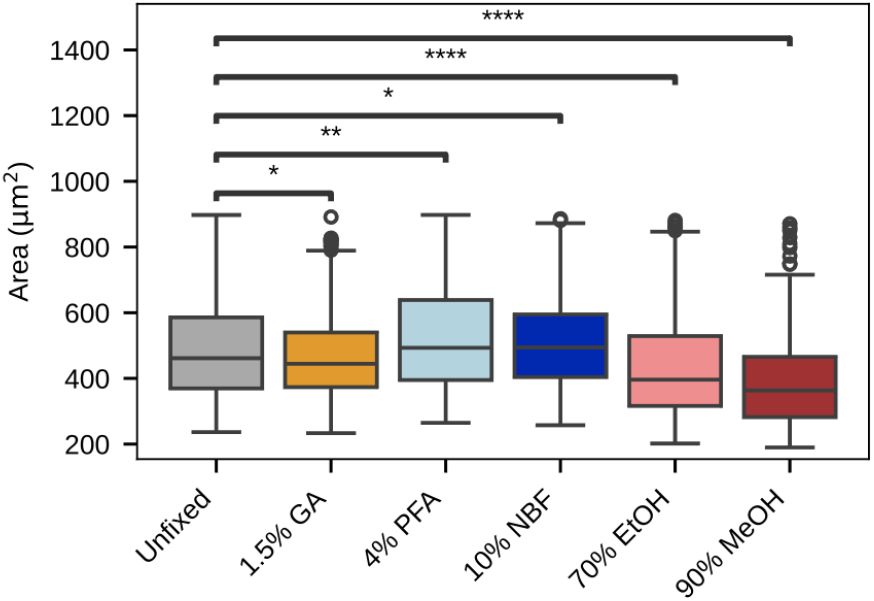
Fixation-induced cell size distributions. Cells are grouped by fixation conditions. *, **, ***, and **** indicate p < 0.05, 0.01, 0.001, and 0.0001 (Welch’s t-test), respectively. Horizontal bars of the boxes represent maximum, 0.75, 0.5, 0.25, minimum, respectively. Outliers are determined by whether the point deviates 1.5 times the interquartile range (0.75 quartile - 0.25 quartile), from either quartile

## Discussion

Here, we used stimulated Raman scattering to investigate the effects of biological fixation on cultured cells to aid in choosing which fixation technique to use in vibrational microscopy. We expand on previous studies characterizing biological fixation techniques in bacteria^14^ and various cell strains^13,17,18^ that use Spontaneous Raman spectroscopy in the fingerprint region, and extend the analysis to the CH region through hyperspectral SRS microscopy. We provide a more comprehensive characterization of fixation-induced changes through high-resolution measurement of chemical and morphological features, combining spectral analysis and quantitative image analyis.

Using a cell model, we described changes in size and chemical composition across different sample preparations, providing guidelines for future stimulated Raman scattering microscopy experiments requiring fixation or designed in tandem with other multimodal techniques. We identified consistent patterns of fixation mechanisms by analyzing a large number of cells and performing imaging over a rangeof days to overcome batch effects. Our results suggest that 4% PFA fixation is the most reliable method to ensure lipid and protein quantity and distributions most consistent with live cell imaging. In agreement with previous work, we observed that formaldehyde-based fixatives may reduce protein concentrations by 5% to 18%, with potential lipid alterations or loss^8,13,17^. Even though a slight size increase was measured, due to swelling following rehydration in PBS, it did not significantly perturb the internal structure of the cell (Fig. 3(a), 4(c,g-h), 5(a)).

GA shows promise as an alternative fixation method for SRS due to the significant signal increase both in lipid and protein, in agreement with previous works^27^, showing minimal spectral shape distortion (Fig. 2(g), 3(a), 4(b,g-h), 5(a)). The signal increase coupled with minimal size variation could suggest a decrease in cell volume, i.e. flattening, resulting in higher cell density^52^, which was not quantified here given the single z-slice images acquired. Further investigation performing 3D imaging could validate this hypothesis, ensuring that the observed effects are not spurious signals introduced by GA binding but are only of morphological nature.

Our observations confirm the considerable disruptive effects of alcohol fixation, not making it a suitable method for SRS as most of the signal is lost, requiring denoising algorithms to recover spectral features^57,58^. Significant structural reorganization was observed (See Fig. 4(e,f), S7), likely due to protein precipitation that leads to a proportional increase (up to 40%, reported in the literature) in concentrations of nucleolar proteins and RNA, due to possible cell shrinkage, and formation of intracellular granules with localized increases in cytoplasmic lipid signal^8^. Moreover, protein denaturation leads to unfolding, exposing or reorienting CH_3_ groups, resulting in aggregation and structural changes that reduce average protein signal^33,46^. These results align with the understanding that cross-linking fixatives best maintain protein conditions similar to live cells^9,17,33^. When coupling an SRS experiment with histological applications or other downstream analyses requiring alcohol fixation, e.g. spatial transcriptomics, significant morphological changes and lipid loss must be taken into account.

Our experiment quantified the effects of fixation on SRS microscopy on a HeLa cell model, informing other cell lines’ possible trends. For a more generalized characterization, the current study could be expanded in scope by testing various human cell models with a more diverse morphology, to further explore the extent of the effects, e.g. the high lipid content of adipocytes makes them particularly susceptible to lipid washout under alcohol fixation conditions. Moreover, this work can suggest possible effects of tissue fixation for SRS imaging, not including morphological factors like extracellular matrix/collagen, overall morphology in 3D, and depth scattering effects coupled with exogenous clearing agents, e.g. paraffin or xylene, used to reduce scattering and allow higher penetration depth for imaging. For these reasons, choosing Raman-compatible clearing methods ensures optimal volumetric chemical imaging with a tenfold increase in penetration depth without distorting the SRS signal^59^. Here, we characterized cellular behaviour focusing on proteins and lipids in the CH region, providing a basis for identifying potential trends when exploring other spectral regions, e.g. fingerprint region, or when using exogenous Raman labels to access the silent region, to study metabolites, or biomolecules^60–62^. The selected concentrations, times, and temperatures are based on a modal sense of the wide variety of methods used in literature. Any variation in these parameters may modulate the fixative-induced effects on SRS, with the patterns described here serving as a reference for interpreting those outcomes.

Our findings provide essential insights into the effects of chemical fixation on cellular lipid and protein profiles, within the CH region of the Raman spectrum. By leveraging label-free, quantitative SRS microscopy, we characterize fixation-induced morpho-molecular changes while minimizing artifacts typically associated with sample preparation. This study represents a pivotal step in advancing SRS as a prime method for biomedical analysis and characterization, ensuring that cells or tissues are observed under the most pristine conditions. We envision the broader application of our findings to refine fixation protocols, enabling more accurate and reproducible Raman-based studies that enhance our understanding of cellular and tissue dynamics in health and disease.

## Acknowledgements

Research reported in this publication was supported by the National Institutes of Health under award number R35GM124916 (A.S.), the Chan Zuckerberg Initiative SDL Award, and the Harvey and Leslie Wagner Foundation. A.S. is a Chan Zuckerberg Biohub Investigator. This work also received funding from the European Union’s Horizon 2020 research and innovation program under grant agreement No. 101016923 (D.P.). D.P. acknowledges financial support by the European Union’s NextGenerationEU Programme with the I-PHOQS Infrastructure [IR0000016, ID D2B8D520, CUP B53C22001750006] “Integrated infrastructure initiative in Photonic and Quantum Sciences”.

## Author Contributions

Conceptualization, A.S., C.C., W.H.; Experimental design, C.C. W.H.; Cell culture and sample preparation, C.C.; SRS imaging, C.C., W.H., B.M.; SRS image analysis, W.H., C.C., B.M.; Writing, C.C., W.H., D.P., A.S., B.M.; Supervision and funding acquisition, A.S., D.P.

