## Supplementary material for "Effects of biological fixation methods on stimulated Raman scattering microscopy signal"

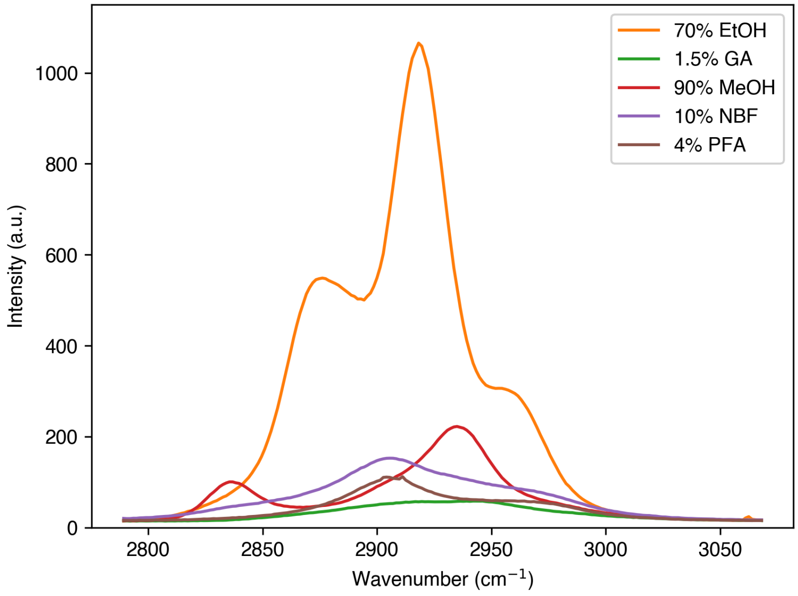


**Fig. S1**: SRL Raman spectrum of the five fixatives in the concentrations employed in the study. See methods for fixative compositions. For this experiment, 50uL of fixative solution is deposited on a coverslip/microscope slide sandwiched configuration and a single-point SRL spectrum is acquired. Laser powers are set to 20/25 mW (Pump/Stokes), wavelengths are the same employed in the study (Pump at 799nm, Stokes at 1040nm) targeting the CH spectral region through spectral focusing. Water peak is outside of the accessible region through spectral focusing at the selected wavelengths.

**Table S2**: Summary of 2920cm$^{-1}$ band (i.e. protein) average intensity distributions across fixatives.

| Fixative | Mean | Std | 25% | 50% | 75% |
|---|---|---|---|---|---|
| 1.5% glutaraldehyde | 591 | 96 | 542 | 604 | 649 |
| 10% formalin | 446 | 65 | 412 | 451 | 489 |
| 4% PFA | 420 | 58 | 379 | 422 | 459 |
| 70% ethanol | 429 | 101 | 352 | 415 | 493 |
| 90% methanol | 388 | 72 | 349 | 398 | 433 |
| unfixed | 511 | 126 | 439 | 541 | 593 |

**Table S3**: Summary of 2850cm$^{-1}$ band (i.e. lipid) average intensity distributions across fixatives.

| fixative | mean | std | 25% | 50% | 75% |
|---|---|---|---|---|---|
| 1.5% glutaraldehyde | 166 | 52 | 144 | 178 | 201 |
| 10% formalin | 133 | 39 | 118 | 137 | 155 |
| 4% PFA | 133 | 42 | 119 | 141 | 162 |
| 70% ethanol | 111 | 43 | 92 | 113 | 134 |
| 90% methanol | 92 | 40 | 79 | 95 | 112 |
| unfixed | 158 | 46 | 132 | 168 | 188 |

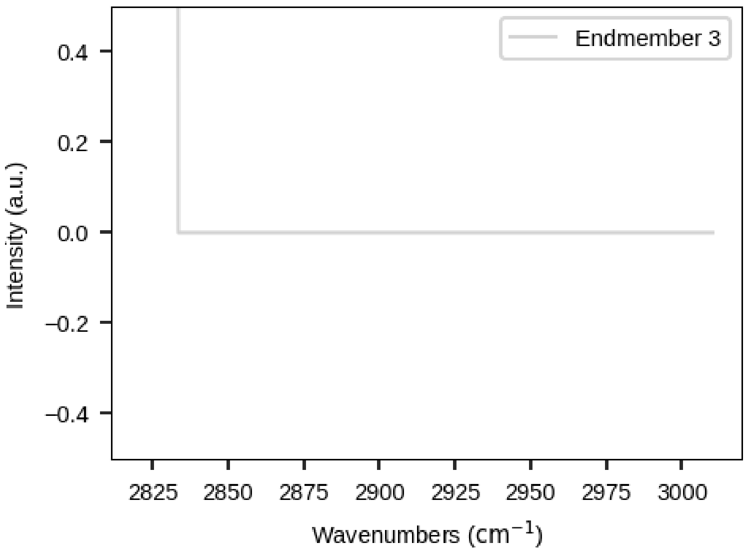


**Fig. S4**: Zoom-in view of the third endmember, contributing mostly noise from wavenumber with the least intensity

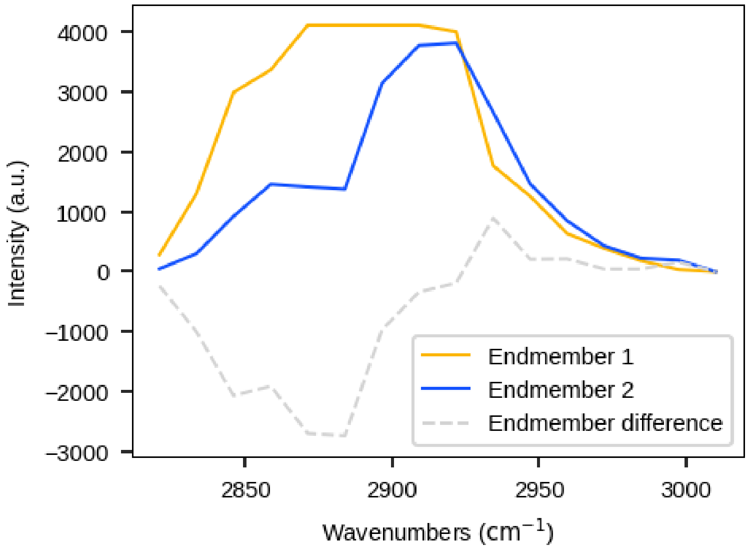


**Fig. S5**: **Spectra of endmembers 1 and 2 extracted using N-FINDR, and their difference.** The selection of 2850 cm$^{-1}$ and 2920 cm$^{-1}$ as representative bands for lipids and proteins, respectively, was guided by the difference spectrum, which highlights regions of maximal variation between the two components. These bands correspond to areas of high contrast and minimal spectral cross-talk, allowing robust discrimination between lipid- and protein-rich regions. Notably, 2850 cm$^{-1}$ and 2920 cm$^{-1}$ are also well-established markers for lipid ($CH_2$ symmetric stretch) and protein ($CH_3$ symmetric stretch) content in the literature.

**Table S6**: Summary of cell size distributions across fixatives. (Units are in μm$^2$)

| Fixative | Mean | Std | 25% | 50% | 75% |
|---|---|---|---|---|---|
| 1.5% glutaraldehyde | 466 | 137 | 373 | 445 | 540 |
| 10% formalin | 508 | 143 | 404 | 494 | 595 |
| 4% PFA | 517 | 156 | 395 | 493 | 639 |
| 70% ethanol | 428 | 153 | 316 | 396 | 529 |
| 90% methanol | 388 | 137 | 282 | 363 | 466 |
| unfixed | 488 | 155 | 370 | 462 | 586 |

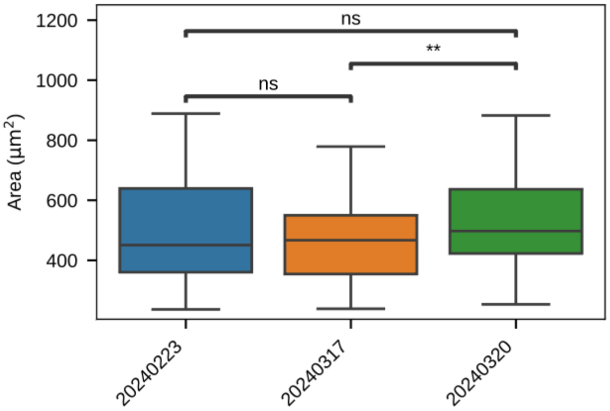


**Fig. S7**: Size distribution of unfixed cells measured on different days. *, **, ***, and **** indicate $p < 0.05$, 0.01, 0.001, and 0.0001 (Welch's t-test), respectively. Horizontal bars of the boxes represent maximum, 0.75, 0.5, 0.25, minimum, respectively. Outliers are determined by whether the point deviates 1.5 times the interquartile range (0.75 quartile - 0.25 quartile), from either quartile. While cell size measurements varied slightly between days, Welch's t-test revealed no statistically significant differences ($p > 0.001$, $p > 0.05$), supporting the conclusion that the measurements originate from the same underlying distribution.

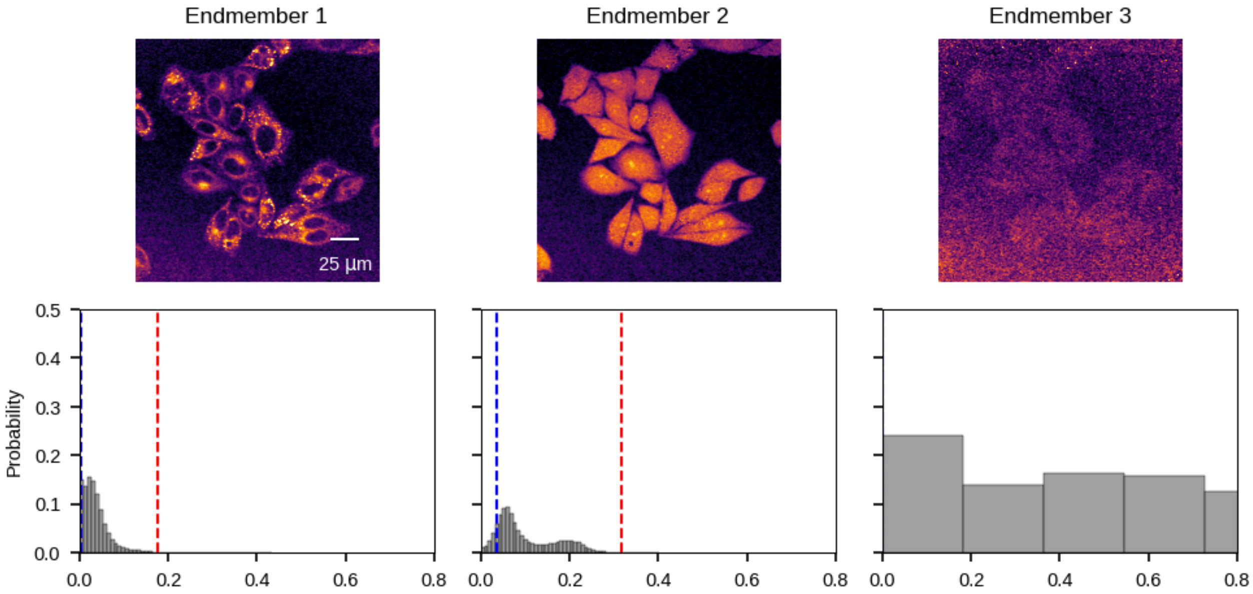


**Fig. S8**: Images of the cells based on non-negative least square reconstruction of endmembers. Bottom plots show the histograms of intensity distributions, as well as the dynamic range for two-color composition.

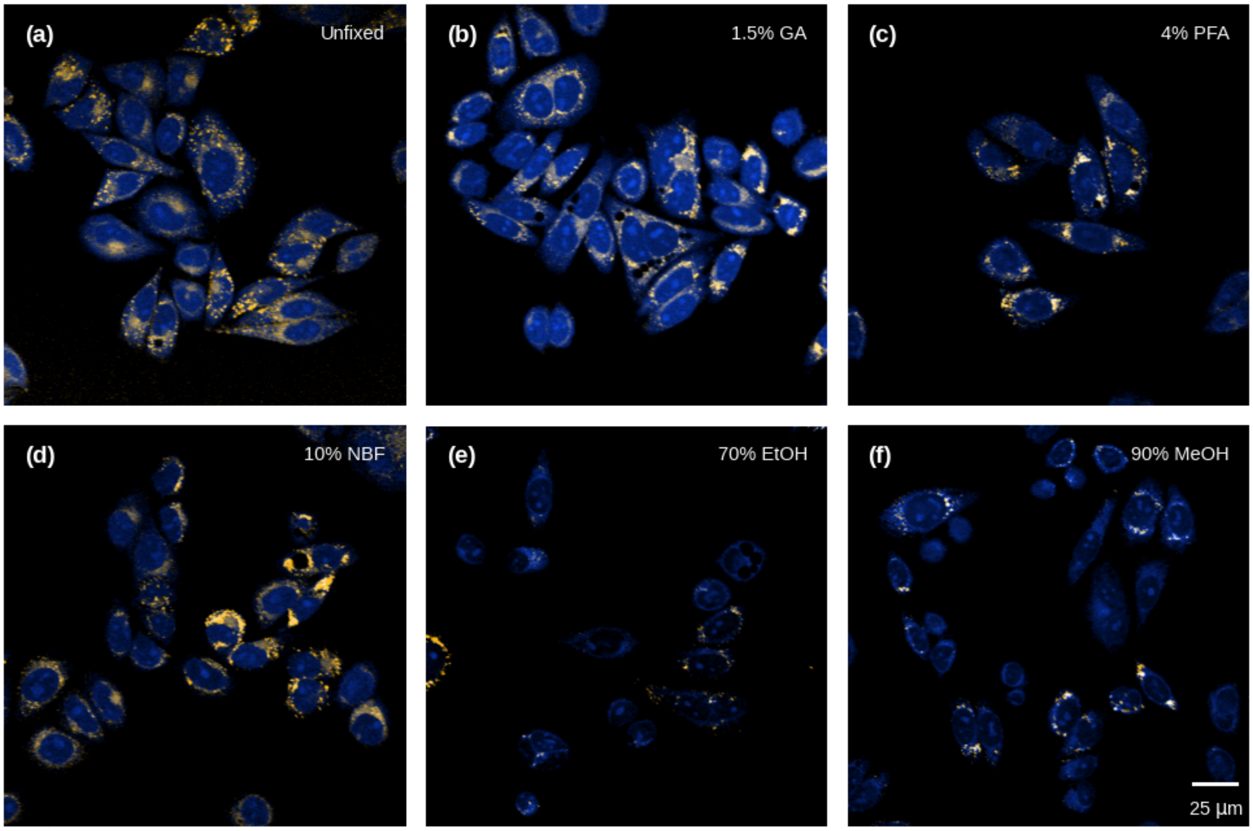


**Fig. S9**: Two-color composition of the reconstructed endmember images. The field of views were the same as shown in Fig. 4.

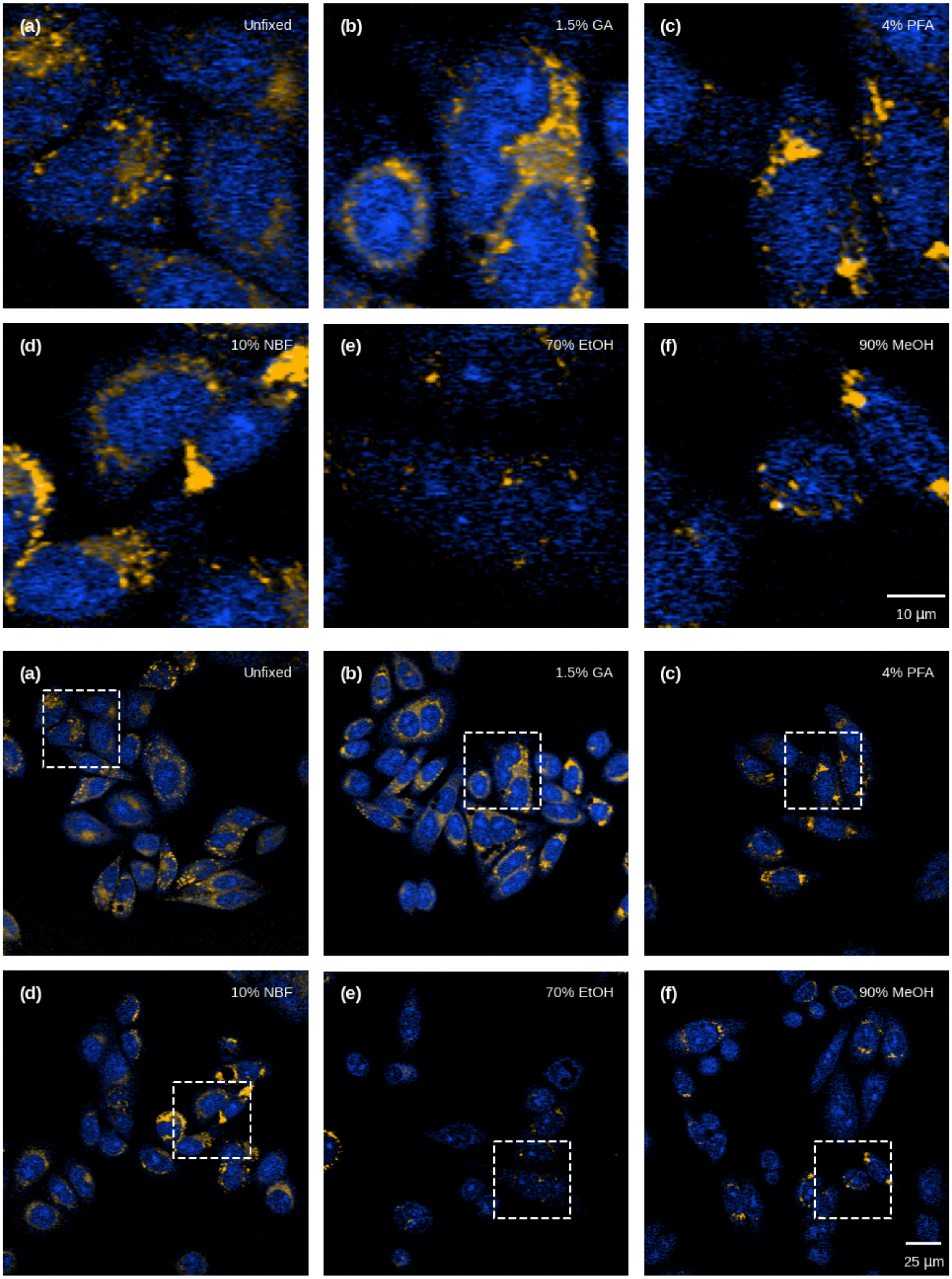


**Fig. S10**: Zoomed-in view of individual cells to highlight changes in nuclear aggregation and their corresponding region in Fig. 4.
